# Anatomy and habitat shape the oxygen sensing machinery of angiosperms

**DOI:** 10.1101/2025.09.17.676756

**Authors:** Ximena Chirinos, Vinay Shukla, Mikel Lavilla-Puerta, Robin Bär, Richard J. Lilley, Angelika Mustroph, Francesco Licausi

## Abstract

Hypoxia sensing via the Cys/Arg branch of the N-degron pathway (Cys-NDP) is central for flooding responses in plants, yet how evolutionary and ecological factors have shaped the core oxygen sensing mechanism remains poorly understood. Leveraging the publication of multiple angiosperm genomes, we systematically analysed known Cys-NDP components in 55 angiosperms spanning aquatic, epiphytic, xerophytic, and mesophytic lineages. We also complemented this survey with hypoxia profiling and transcriptomic analyses in a selected panel of plants. This comparative effort revealed variation in Cys-NDP components, with Plant Cysteine Oxidases (PCOs) and group VII Ethylene Response Factors (ERFVIIs) emerging as major sources of diversification. Aquatic monocots displayed complete loss of A-type PCOs and dramatic expansion of a novel clade of ERFVIIs (HRE_aqua_), frequently accompanied by loss or modification of the Cys-degron, uncoupling them from oxygen-dependent turnover. By contrast, xerophytes and epiphytes retained core Cys-NDP elements but showed shared hypoxia-induced gene expression, suggesting endogenous developmental or metabolic pressures for pathway conservation in habitats with limited flooding risk. Across all species, we identified a conserved transcriptional core of 11 orthogroups, including fermentation enzymes and regulatory factors, highlighting the early recruitment of these genes to hypoxia responses. Functional assays confirmed contributions of conserved MYB and LBD transcription factors to hypoxia tolerance in *Arabidopsis*. Together, our results demonstrate that both habitat and anatomy influence the evolution and deployment of oxygen-sensing networks in angiosperms. While persistent submergence promoted diversification of ERFVIIs and PCOs, retention of the core pathway across lineages points to fundamental roles in coping with endogenous oxygen gradients and fluctuations.

## Introduction

Molecular oxygen (O_2_) is an essential substrate for numerous metabolic reactions, most notably oxidative phosphorylation, which enables chemical energy storage in the form of ATP. Because of this key role, plants have evolved mechanisms to sense oxygen limitation (hypoxia) and adjust their metabolism and development accordingly. While information about the strategies adopted by green algae and bryophytes is limited, recent work has shown that all vascular plants detect oxygen through the Cys/Arg branch of the N-degron pathway (Cys-NDP) for protein degradation^1^.

This proteolytic mechanism links cellular oxygen availability to the stability of proteins bearing an N-terminal cysteine (N-Cys), which are generated either through removal of the initiator methionine by methionine aminopeptidases (MetAPs) or through cleavage by endoproteases^2,3^. Proteins are targeted for degradation by Plant Cysteine Oxidases (PCOs), which oxidize N-Cys to sulphinic or sulphonic acid^4,5^. The modified residue is subsequently arginylated by Arginine-tRNA transferase (ATE), generating a substrate for polyubiquitination by the giant N-recognins PRT6 and BIG^6^. These enzymes target the modified proteins for proteasomal degradation^6^. Although the conservation of N-Cys residues across protein families and the pleiotropic phenotypes of NDP mutants suggest that many substrates exist, only three have been experimentally validated to date: the group VII Ethylene Response Factors (ERFVIIs), Vernalization 2 (VRN2), and Little Zipper 2 (ZPR2)^7–10^.

Among these, ERFVIIs are conserved in vascular plants and mediate transcriptional responses to hypoxia, whereas VRN2 and ZPR2 are known Cys-NDP substrates only in angiosperms^1^. ERFVIIs act within complex feedback networks, inducing the expression of other ERFVIIs, PCOs, and interacting partners that fine-tune their activity^4,11,12^. Initially identified as key regulators of metabolic adaptation to hypoxia^11,13^, ERFVIIs have since been implicated in diverse physiological and developmental processes, including seed germination and dormancy, seedling establishment, stomatal regulation, root growth, and general stress responses^14^.

Most characterization of this oxygen-sensing pathway has been carried out in model angiosperms such as *Arabidopsis thaliana, Oryza sativa*, and *Medicago sativa*^15–17^. These species frequently experience fluctuations in oxygen availability in their natural habitats, due either to restricted environmental O_2_ levels (e.g., during submergence or waterlogging) or to internal constraints such as high respiratory demand relative to diffusion, as in shoot apical meristems. A recent report found that ERFVII and PCO genes were expanded in the genomes of four seagrass species due to genome multiplication, a change hypothesized to facilitate adaptation to marine environments^18^. By contrast, little is known about species that are less likely to encounter hypoxia due to their anatomy or ecological niches. This gap is particularly intriguing in angiosperms, given their extraordinary diversity and global distribution.

Although molecular dating methods disagree on whether angiosperms originated early or late in the Mesozoic, phylogenetic analyses consistently reveal multiple waves of rapid diversification. The first major expansion, the Angiosperm Terrestrial Revolution, occurred across Magnolids, Monocots, and Eudicots at different points in the Cretaceous, followed by slower but steady diversification that gave rise to most extant families^19,20^. Later surges, especially in Eudicots, may have been driven by insect–angiosperm coevolution and by climatic cooling during the Cenozoic^20^. Interestingly, from an hypoxia perspective, some terrestrial angiosperms independently colonised aquatic habitats through key adaptations^18^.

Intrigued by the conservation of the oxygen-sensing Cys-NDP branch and its involvement in diverse molecular and physiological processes, we set out to examine the relationship between anatomy, habitat, and the composition of oxygen-sensing enzymes and substrates. In particular, we examined whether exogenous (environmental) and endogenous (developmental or anatomical) drivers of hypoxia have shaped the diversity, function, and regulatory scope of this pathway in angiosperms.

## Results

We explored the pool of available angiosperm genomes and selected 54 species belonging to Nympheaeles (1), Ceratophyllales (1), monocot (24), and eudicot (28) clades, representing extremes in their likelihood of experiencing submergence-related hypoxia during the life cycle (**Supplementary Table 1**). Macrophytes (aquatic plants, 16 species) are constantly submerged, whereas xerophytes (desert plants, 7 species) and epiphytes (aerial plants, 7 species) are unlikely to encounter submergence in their natural habitats. They are further characterized by diverse morphologies, such as leaf shape, stem thickness, types of anchorage organs, and growth rates, which may generate different oxygen gradients and fluctuations within their organs. As a control group, we also included 24 mesophyte species that are likely to experience only occasional hypoxia throughout their lifecycle. The uneven distribution of clades across the three hypoxia-related habitats, with most macrophytes and epiphytes being monocots, while xerophytes were predominantly eudicots, was due to genome availability and did not necessarily reflect clade-specific adaptive capacity. For example, no genome of an epiphytic eudicot was available at the time of this analysis.

Applying the Orthofinder algorithm^21^, we searched this pool of selected genomes for orthologous genes coding for Cys-NDP enzymes and substrates. We first focused on the copy numbers of each orthogroup (OG) of interest (**Fig. 1a**). Our survey confirmed the previously observed modest conservation of ZPR2 proteins as substrates of the Cys-NDP^9^ across all angiosperm species considered. This analysis also revealed that these are absent or limited to one single instance in macrophytes. VRN2 was not conserved as Cys-NDP substrate in the eudicot family Amaranthaceae (**Fig. 1a**).

**Fig. 1:**
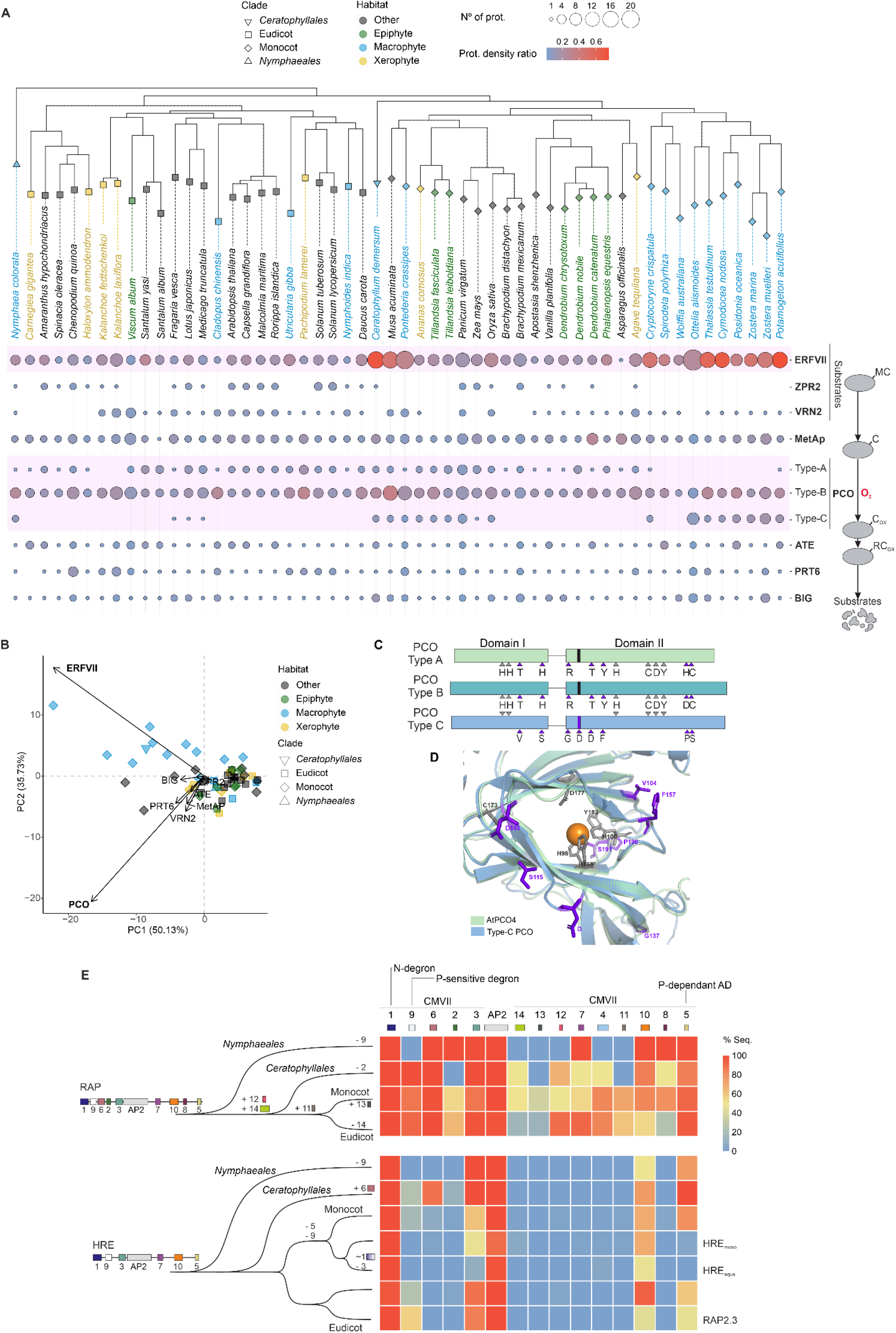
The Cys-NDP across angiosperms that colonized habitats with opposite submergence probability. **a.** Bubble plot depicting the distribution of known Cys-NDP enzymes and their substrates across macrophytic, epiphytic, xerophytic and mesophytic angiosperms. **b**. PCA assessment of the two major drivers of Cys-NDP differentiation across selected angiosperms. **c-d**. Comparison of A-type and C-type PCOs highlighting in purple the amino acid substitutions on a linear representation (c) and on a 3D overlay between the predicted structure of a C-type PCO consensus and the experimentally determined *Arabidopsis thaliana* PCO4 (PDB 6S7E) (d). **e**. Evolution of RAP and HRE proteins in angiosperms as reconstructed from the occurrence of conserved motifs.

Principal component analysis of the selected species, considering the Cys-NDP components, identified ERFVIIs and PCOs as the major drivers of diversity (**Fig. 1b**). Indeed, PCOs and ERFVIIs represented the most populated OGs among Cys-NDP enzymes and substrates, respectively, suggesting an evolutionary preference for modulating hypoxia responses through these genes (**Supplementary Table 1**). While we could not observe specific trends for epiphytes and xerophytes, macrophytes showed reduced numbers of A-type, constitutive, PCOs, with almost-complete disappearance of this OG in aquatic monocots (**Fig. 1a**).

In addition to the previously reported A-type and B-type clusters^22^, Orthofinder identified a third PCO OG, which we termed C-type, largely comprising monocot sequences. This OG also contained one sequence each from Ceratophyllales and Nymphaeales, suggesting an ancient origin (**Fig. 1a**). C-type PCOs were found in only three eudicots, which could be explained by multiple independent losses in this clade (**Fig. 1a**). This OG was remarkably expanded in aquatic monocots, which also possessed significantly more ERFVIIs—unlike aquatic eudicots (**Fig. 1a**). Interestingly, species from the aquatic Ceratophyllales shared this feature, whereas Nymphaeales, which mainly grow at the water surface, did not. Across monocot macrophytes, the ERFVII-to-PCO ratio was consistently higher (**Fig. 1a, Supplementary Fig.1a**). We therefore hypothesize that colonization of aquatic environments, characterized by frequent hypoxia, has reshaped the architecture and stoichiometry of the Cys-NDP.

Intrigued by the uneven distribution of *PCO* and *ERVII* genes, we next looked at their diversity in terms of encoded amino acid sequences. PCOs showed strong conservation, with invariant residues required for substrate coordination and catalytic activity (**Supplementary Fig. 2a**). The predicted structure of a consensus C-type PCO sequence largely overlapped with those obtained for Arabidopsis PCO2 and PCO4^23^ (**Fig. 1c, Supplementary Fig. 2b**). They retain the distinctive cupin β-barrel preceded by two α-helices that forms a wide cavity sufficient to accommodate peptide substrates^24^ (**Supplementary Fig. 2b-c**). In this, we could identify three conserved His residues, which in PCO2 and 4 coordinate Fe(II). C-type PCOs also retained Glu and Tyr, which have been proposed to contribute to substrate positioning, and Cys173 (according to PCO4 numbering (**Fig. 1c-d, Supplementary Fig. 2a-b**). C-type PCOs could be distinguished from A- and B-type PCOs by the conversions Thr104Val, Arg136Gly, Tyr157Phe, the addition of an additional Asp/Glu in position 143 and lack of conservation at positions 115, 148 and 190-201 (**Fig. 1c-d, Supplementary Fig. 2a**). Additionally, we often found a poly Asp/Glu tail of variable length at the C-termini of C-type PCOs. Surprisingly, PCO-C sequences from aquatic monocots carried a conserved potential N-Cys-degron (**Supplementary Fig. 2d**). Based on these predictions, we speculate that C-type PCOs participate to the Cys-NDP as A- and B-type PCOs do, although with kinetics that might accommodate substrate (de)stabilisation at slow oxygen dynamics and steep gradients expected for aquatic plant species.

**Fig. 2:**
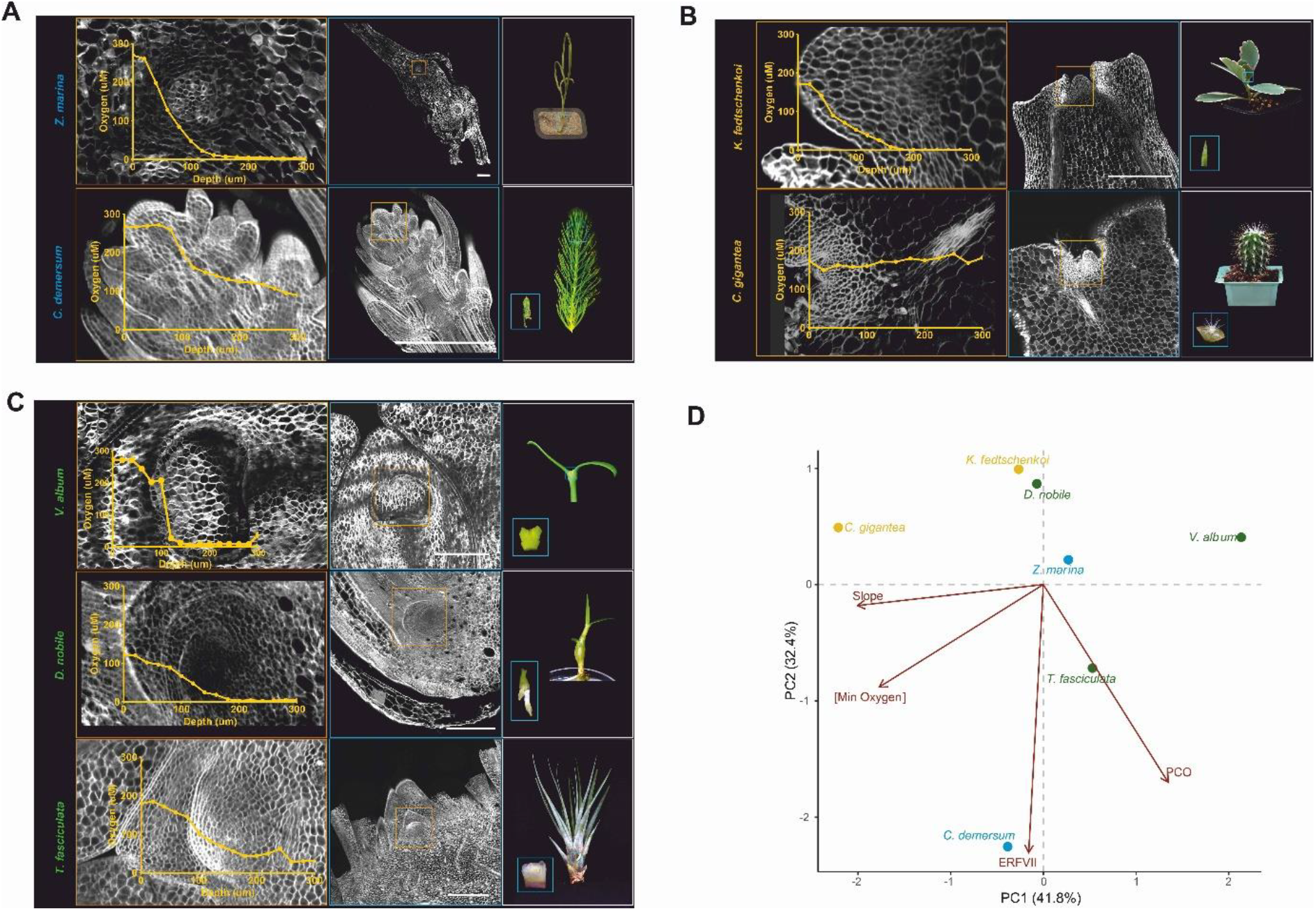
Oxygen profiles of SAM across a selected panel of macrophytes, epiphytes and xerophytes. Oxygen concentration though the SAM in the apical-to-basal direction overlapped to SAM longitudinal section of *Z. marina* and *C. demersum* (macrophytes) in **a**, *V. album, D. nobile* and *T. fasciculata* (epiphytes) in **b** and C. gigantea and *K. fedshikenkoi* (xerophytes) in **c**. Progressive zoom out show photographs of SAM location in the plant. **d**. PCA assessment of O_2_-minima, O_2_-slopes and PCO and ERFVII gene numbers across selected angiosperms.

In contrast to PCOs, ERFVIIs exhibited high variability outside of the AP2 DNA-binding domain, consistent with relaxed functional constraints. We conducted an unbiased motif search in ERFVII sequences, recovering sequences previously described in Nakano et al.(2006)^25^ but also identifying five more (CMVII-9-14). Their presence/absence patterns allowed us to cluster angiosperm ERFVIIs into four main clades (**Fig. 1e; Supplementary Fig. 3**). Two major OGs included orthologs of the constitutively expressed RAP2.2 and RAP2.12 (RAPs) and of the hypoxia-inducible ERFs (HREs) of *A. thaliana*, respectively^11^. Both share the AP2 DNA-binding domain and four additional motifs (CMVII-1, 3, 5 and 10), including the N-Cys-degron and phosphorylation-dependent transcription activation domain (AD). Given their ubiquitous conservation, we considered them as unvaried descendants of two separate ancestral genes already present in the last common ancestor of vascular plants^1^ (**Fig. 1e**).

**Fig. 3:**
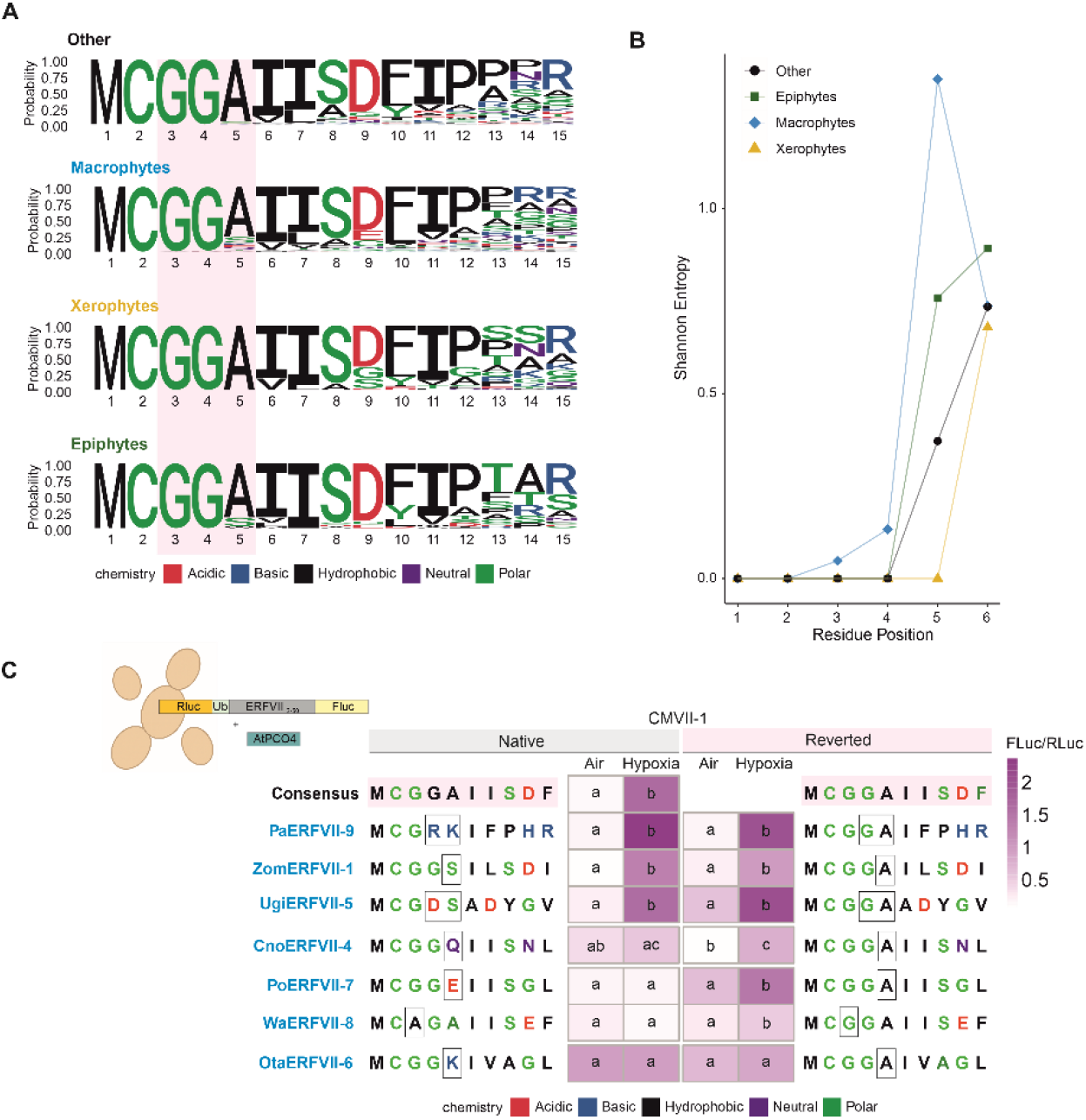
Cys-degron variation in macrophyte ERFVIIs. **a.** Logos depicting amino acid occurrences at positions 1-15 in ERFVIIs from the macrophyte, xerophyte and epiphyte species analysed in Fig. 1a. Proteins that lost the Cys-degron were excluded from this analysis. **b**. Sequence conservation and variability at positions 1-6 in the ERFVII considered in a. **c**. Protein stability measured by yeast DLOR assay for a selected panel of seven HRE_aqua_ showing variation in the conserved CMVII-1 (left) and their mutant variants where the consensus sequence was restored (right). Letters denote statistically significant groups with different stability (p≤0.05, one-way ANOVA followed by Tukey post-hoc test).

The RAP clade shares CMVII-2, 3, 6, 7 and 8 and in Mesangiospermae acquired CMVII-12 and 14. CMVII-9, which corresponds to a phosphorylation-sensitive degron^26^, was likely present in the last common ERFVII ancestor and then lost multiple times (**Fig. 1e, Supplementary Table 2**). In RAPs this happened only in Nimphaeales, whereas in HREs the evolutionary pressure to retain it was significantly lower (**Fig. 1e, Supplementary Table 2**), potentially to evade Ca^2+^-dependent regulation^26^. Additional motifs were acquired and lost in monocots and eudicots. In contrast to the RAPs, the HRE clade duplicated and diverged multiple times. In Ceratophyllales, it acquired the CMVII-6 and in monocots, it split twice to generate three clades: HRE, HRE_mono_ and a macrophyte-specific one, which we termed HRE_aqua._ In eudicots, HREs duplicated to give rise to a separate clade that includes the Arabidopsis RAP2.3, (**Fig. 1e, Supplementary Table 2**).

In the HRE clade, we found (29) ERFVIIs which lost the CMVII-1, corresponding to this group’s distinctive N-degron (MCGGAI), and therefore cannot be regulated by O_2_ availability (**Supplementary Fig.1a, Supplementary Table 1**). Discharge from the Cys-NDP occurred independently across all HREs, but not in RAPs. Noticeably, these events were over-represented in monocot macrophytes (**Supplementary Fig.1d, Supplementary Table 1)**. Altogether, these observations support the view of ERFVII and PCO as means to accommodate monocot’s adaptation to aquatic habitats.

Since we mainly observed an effect of the aquatic habitat on ERFVIIs, whereas limited exposure to submergence did not influence Cys-degron distribution, we tested whether anatomical features leading to O_2_ restriction could bias gene number. We focused on the shoot apical meristem (SAM), a crucial tissue for both vegetative and reproductive growth, which is conserved across all phylogenetic groups included in our metagenomic analysis.

From our list of 30 angiosperms adapted to contrasting environments, we selected and grew seven species: aquatic (*Zostera marina, Ceratophyllum demersum*), arid (*Kalanchoë fedtschenkoi, Carnegiea gigantea*), and epiphytic (*Viscum album, Dendrobium nobile, Tillandsia fasciculata*). For each of this species, we measured O_2_ levels in SAM tissues using a microscopic Clark electrode. All SAMs except that of *C. gigantea* showed a decline in oxygen availability toward the core, consistent with observations in *Arabidopsis* and tomato^9^ (**Fig. 2a**). This observation extends the concept of SAM hypoxia to the monocot clade. We hypothesized that the relatively high oxygenation of the *C. gigantea* SAM is due to the slow production of ribs (orthostichies) compared with the fast plastochron reported in the other species^27,28^. In contrast, photosynthetically active organs (leaves and stems) in the seven species considered did not display O_2_ gradients (**Supplementary Fig. 4**). Finally, a principal component analysis (PCA) did not reveal concordant trends between O_2_ gradients in the SAMs and the copy numbers of PCOs and ERFVIIs (**Fig. 2b**). We therefore concluded that habitat, and especially aquatic environments, rather than anatomy drove variations in the assortment of Cys-NDP components involved in O_2_ sensing.

**Fig. 4.**
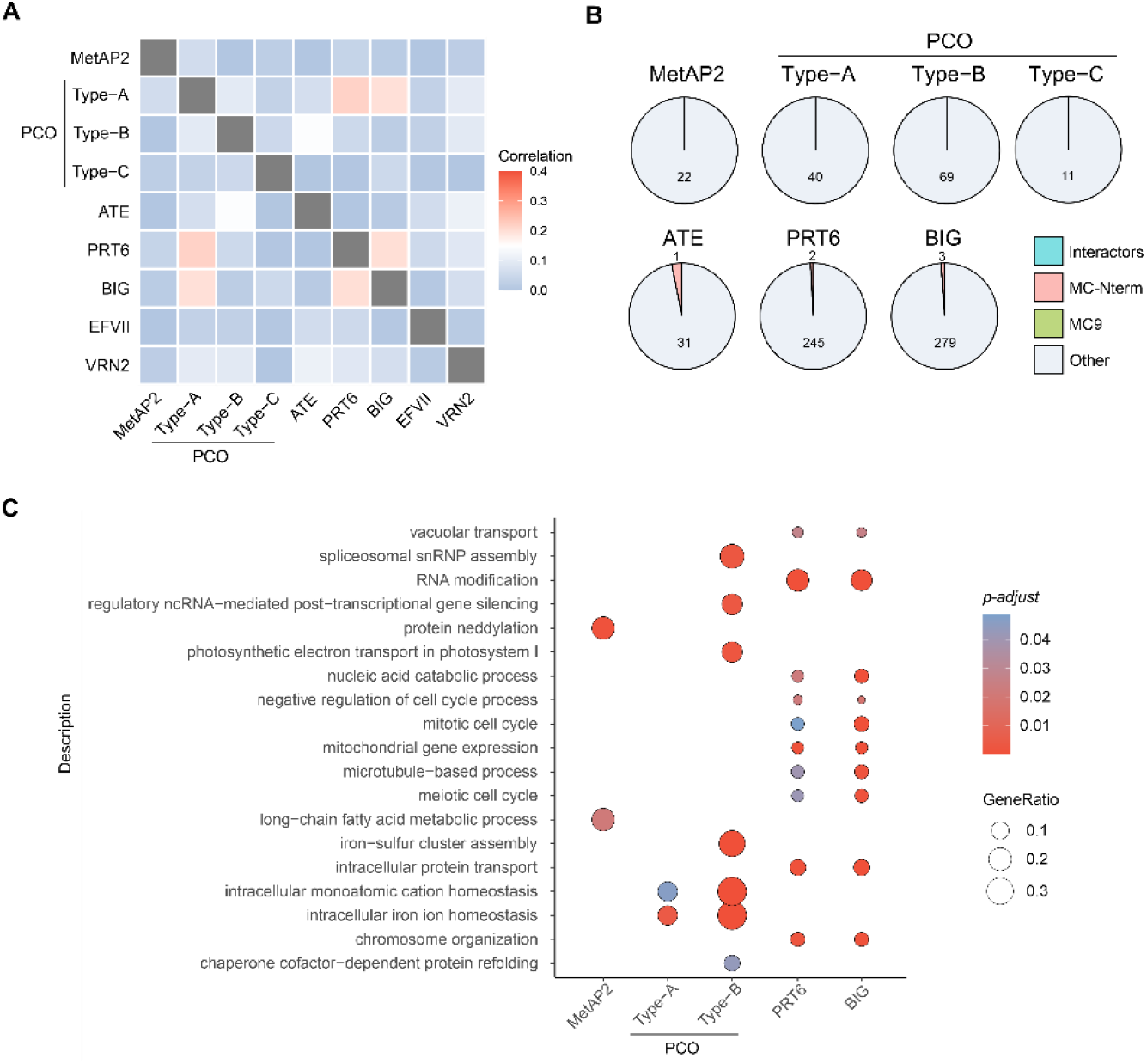
Evolutionary rate covariation (ERC) of Cys-NDP components in angiosperms. **a.** Sequence evolution correlation between known Cys-NDP enzymes and substrates. ZPR2 was excluded because ERCnet could not distinguish it from ZPR1. **b**. Pie chart showing the overlap between predicted Cys-NDP enzyme substrates, either because MC-proteins or substrates of Metacaspase 9 in Arabidopsis, or interactors and the proteins showing significant ERC with Cys-NDP enzymes. **c**. Gene Ontology term enrichment for proteins showing significant ERC with Cys-NDP enzymes.

When focusing on HREs_mono_ of aquatic monocots, we noticed divergence in the otherwise strongly conserved CMVII-1 motif, which corresponds to the Cys-degron_8_. While in other instances described above the whole sequence was absent (**Supplementary Table 1**), here most of the CMVII-1, including the Cys2, was retained, although substitutions occurred in position 3, 4, or 5. Later positions were equally variable in xerophytes, epiphytes and macrophytes (**Fig. 2a and b**). To test whether these substitutions affected ERFVII stability via the N-degron pathway, we used an established heterologous assay in *Saccharomyces cerevisiae*_29_. We compared the stability of FLuc-fused CMVII-1 variants under normoxia and hypoxia, both in their native form and after restoring the sequence to the consensus.

The tested sequences displayed three types of behaviour. First, three variants with charged residues at positions 4 or 5 showed hypoxia-dependent stabilization in both their native and consensus-restored forms (**Fig. 3a**). Second, *CnoERFVII-4*, carrying a Gln at position 5 (polar but uncharged), exhibited limited destabilization that was not significantly relieved in hypoxia. Substitution of Gln5 with Ala restored hypoxia-dependent stabilization (**Fig. 3a**). Similarly, *PoERFVII-7*, with a Glu at position 5 (polar, negatively charged), showed constitutive destabilization regardless of O_2_ availability, but reversion to Ala re-established hypoxia-dependent stabilization (**Fig. 3a**). A comparable pattern was also observed when Ala replaced Gly at position 3. Finally, *OtaERFVII-6*, carrying a Lys at position 5 (polar, positively charged), was constitutively stable, and oxygen-dependent degradation could not be restored by the Lys5Ala substitution (**Fig. 3a**).

We conclude that mutations in the CMVII-1 domain influence susceptibility to the PCO-dependent N-degron pathway, although in some cases additional residues outside conserved regions also contribute to regulation. By contrast, members of the constitutive RAP clade possess an additional domain that may help define them as Cys-NDP substrates.

The high number of enzymatic components and substrates sequences of this O_2_-sensing pathway allowed us to determine their evolutionary rate covariation (ERC) in angiosperms. Using the ERCnet algorithm^30^, we detected correlations between the evolutionary rates of constitutively expressed PCOs and PRT6/BIG, as well as between BIG and PRT6 (**Fig. 4a, Supplementary Table 2**). Since BIG and PRT6 are both large proteins that have been proposed to interact or cooperate, it is unsurprising that co-occurring mutations may help them adapt to recruit the same substrates or partners. More intriguing was the correlation observed for A- and B-type PCOs, which may reflect a requirement to bind or interact with similar N-degrons substrates (**Fig. 4a**). Although below our chosen significance threshold of R^2^ = 0.2, a weaker correlation was also detected between hypoxia-inducible PCOs and ATE substrates (**Fig. 4a**). The Cys-NDP substrate VRN2 further showed correlations with all N-terminal processing enzymes except MetAPs substrates (**Fig. 4a**).

We also assessed ERC between N-terminal processing enzymes and the entire angiosperm metaproteome. This analysis identified ample variation in the number of correlating proteins, which spanned one order of magnitude between MetAP and large ubiquitin E3 ligases (**Fig. 4b**). Within these datasets, we searched for known interactors, proteins exposing an N-Cys due to MetAP activity at Met1^3^, or as a result of metacaspase-9 (MC9)-mediated endoproteolysis^31^. Among the evolutionary correlating proteins, we detected only a few Cys2 proteins but no known interactors or MC9 substrates (**Fig. 4b**). A gene ontology (GO)-enrichment analysis showed significant over-representation of proteins involved in housekeeping processes for PRT6 and BIG, including cytoskeleton dynamics and the cell cycle. B-type PCOs co-evolved with proteins involved in RNA processing, protein folding and iron homeostasis (**Fig. 4c**). Only this latter feature was shared with A-type PCOs. MetAPs instead specifically co-evolved with proteins involved in neddylation and fatty acid metabolism (**Fig. 4c**).

In summary, the ERC analysis provided interesting insights in the participation of NDP enzymes, including PCOs, to biological processes beyond oxygen sensing, but was not useful in the search of novel substrates. It is likely that proteins that display high sequence plasticity, such as transcription factors or immune receptors, are not effectively captured by this method.

We next tested how the evolution of the Cys-NDP affected the transcriptional response to hypoxia in species that colonize habitats with different flooding probability. To this end, we performed RNA sequencing of the seven species previously selected to study the impact of anatomy on hypoxia gradients. For each species, we sampled vegetative, photosynthetically active organs from plants grown under normoxic conditions and exposed them to hypoxia (1% O_2_, 8 h). For aquatic species, normoxic conditions were maintained by bubbling air into the water, and dissolved O2 concentrations were monitored (**Supplementary Fig. 5**).

**Fig. 5.**
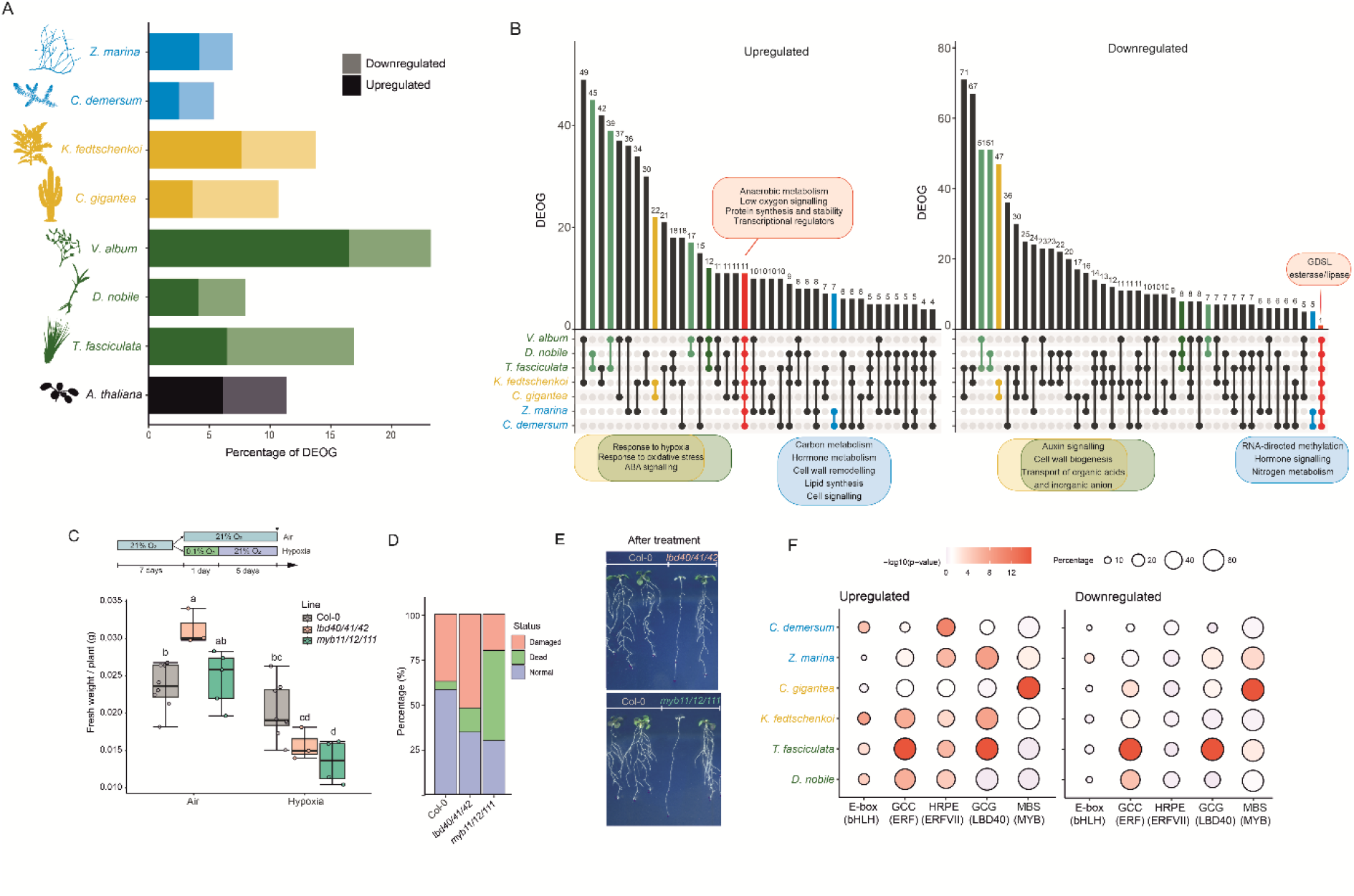
Transcriptional response to hypoxia in selected macrophytes, xerophytes and epiphytes. **a.** Relative number of significantly up- and down regulated OGs (DEOGs) in *Z. marina* and *C. demersum* (macrophytes, blue), *V. album, D. nobile* and *T. fasciculata* (epiphytes, green) and *C. gigantea* and *K. fedshikenkoi* (xerophytes, ochre). *A. thaliana* data from Dalle Carbonare et al. (2025) are included to provide a mesophye control. Significantly differently expressed OGs are selected as those including at least one gene with log_2_FC > |1| (adj. p ≤ 0.05). The number of DEOG was normalized by the total number of common OGs in the species genome. **b**. Shared hypoxia up-regulated (left) or down-regulated (right) DEOGs among the species listed in a. Macrophytes are labelled in blue, epiphytes in green and xerophytes in ochre. Boxes list molecular functions of shared OGs. **c-e**. Hypoxia tolerance of Arabidopsis *myb11/111/112* and *lbd40/41/42* mutants compared to wild type Col-0. Biomass accumulation 5 days after the 24 h-long normoxic (control) and hypoxic treatment is shown in c. Percentage of phenotypes (dead, alive and damaged) measured after the hypoxic treatment is shown in d. An example of wildtype and mutant plants 5 days after the hypoxic treatment is shown in e. **f**. Bubble plot showing the occurrence, in the promoter of hypoxia responsive genes, of DNA elements recognised by transcription factors identified as ubiquitously induced in angiosperms.

The number of differentially expressed genes (DEGs) varied markedly among species (**Supplementary Table 3, Supplementary Fig. 6a**). To enable cross-species comparisons, we referred to differentially expressed OGs (DEOGs). OGs were defined as up-or down-regulated when at least one member gene showed significant induction or repression (log_2_FC > |1|, adj. p ≤ 0.05). Macrophytes exhibited fewer differentially DEOGs than xerophytes and epiphytes, both in terms of induction and repression. The ratio of induced to repressed OGs did not correlate with habitat of origin, but instead appeared species-specific, with the epiphyte *Viscum album* (mistletoe) displaying the highest number of hypoxia-induced Ogs (**Fig. 5a**). We also observed a lower magnitude of gene induction or repression in macrophytes (**Supplementary Fig. 6a**). We interpreted this pattern as reflecting a constitutive habituation of macrophytes to O_2_-limited conditions throughout their life cycle. Conversely, species less likely to encounter hypoxia appeared to mount a stronger transcriptional response when exposed to it. Indeed, *Arabidopsis thaliana* (Col-0), which can occasionally experience flooding-induced hypoxia in its natural habitat, showed an intermediate response.

To identify ubiquitous responses, we searched for OGs that were induced or repressed in at least six of the seven examined angiosperms, and manually inspected whether, in the remaining species, they showed differential expression (log_2_FC > |1|) without reaching statistical significance (adj. p ≤ 0.05). Across all species, we identified a core of 11 upregulated OGs and a single downregulated OG, the latter encoding a GDSL esterase/lipase protein (**Fig. 5b**). The conserved upregulated OGs included well-established hypoxia-responsive genes involved in anaerobic metabolism (Sucrose Synthase, Alcohol Dehydrogenase, Pyruvate Decarboxylase), O_2_ sensing and signal transduction (including B-type PCOs and a CBL-interacting Ser/Thr kinase), and proteins implicated in translation and protein stability (penta- and tetra-tricopeptide repeat proteins, and ADP-ribosyltransferases) (**Fig. 5b**). Several transcriptional regulators were also part of this group, including HRE-type ERFVIIs, the trihelix attenuator HRA1, Lateral Organ Boundary domain (LBD) proteins, class Ia basic Helix-Loop-Helix (bHLH) proteins, and MYB transcription factors (**Fig. 5b**).

When looking at responses conserved across species with similar likelihood to be flooded, epiphytes and xerophytes shared many conserved DEOGs, which were enriched in responses to oxidative stress and ABA signalling genes among up-regulated genes and auxin signalling, cell wall biogenesis and transport of organic acids and inorganic anion among down-regulated OGs (**Fig. 5b, Supplementary Table 4**). Macrophytes instead shared activation and repression of 7 and 5 OGs, respectively (**Fig. 5b, Supplementary Table 4**). The first are likely involved in carbon metabolism, hormone metabolism, cell wall remodelling, lipid synthesis and cell signalling. The down-regulated ones relate to RNA-directed methylation, hormone signalling and nitrogen metabolism. The newly identified C-type PCO clade (**Fig. 1a**) was conservatively induced by hypoxia in these aquatic species (**Supplementary Table 4**).

Three of the conserved hypoxia-induced OGs encoding transcription factors had not previously been functionally characterized in relation to hypoxia tolerance: R2R3-MYBs^32^, class Ia bHLHs^33^, and class II LBDs^34^. We confirmed that the corresponding Arabidopsis genes are induced by hypoxia and tested their relative mutants for low O_2_ tolerance (**Supplementary Fig. 7**). When seedlings of these mutants were exposed to a strict hypoxia treatment (0.1% O_2_ for 24 h), they exhibited a significant reduction in survival and biomass accumulation compared with the wild ecotype Columbia-0 (**Fig. 5c**). These results indicate that the corresponding transcription factors contribute to hypoxia tolerance in Arabidopsis. Given their conserved induction across all species analysed, this role is likely conserved more broadly among angiosperms.

We finally examined the promoters (2 kb upstream and 0.5 kb downstream of the transcription start site) of DEGs for known binding motifs of transcription factors encoded by conserved upregulated genes: the G-box for class Ia bHLHs^33^, the generic GCC-box and the specific HRPE motif for ERFVIIs^35^, the GCG-box for LBDs^36^, and the MYB-binding site (MBS)^37^. *V. album* was excluded from this analysis because its genome sequence was not available at the time of this analysis. We observed significant enrichment of HRPEs in the promoters of hypoxia-induced genes in five of the six species, supporting a conserved role of ERFVIIs as activators of the hypoxia response^1^ (**Fig. 5d**). Interestingly, *C. gigantea* showed no HRPE enrichment in either promoter set, suggesting a potentially diminished role of ERFVIIs in this species (**Fig. 5d**). No habitat-specific bias was detected for the other elements, which were enriched in at least one representative species of each habitat type, with the exception of MYB-binding sites, which were not significantly enriched in epiphytes (**Fig. 5c**). Among promoters of the repressed genes, the E-box and the HRPE were not enriched, whereas the GCC-box, GCG-box, and MBS were overrepresented in at least three of the six species analysed, suggesting conserved separation in the role as gene inducers and repressors among transcription factors.

## Discussion

The extraordinary anatomic diversity and habitat plasticity of angiosperms triggered our curiosity to test how the Cys-NDP oxygen sensing mechanism, which intersects many developmental and environmental response pathways, has diversified alongside plant architecture and ecological niches. Specifically, this study addressed whether the likelihood of experiencing hypoxia due to environmental or developmental factors acts as an evolutionary driver of hypoxia signaling machinery in angiosperms. Our results show that both factors influence the diversity, size, and physiological relevance of the Cys-NDP in an interdependent manner. The analysis revealed that diversification mainly occurred through duplication of PCOs and ERFVIIs. In contrast, as previously reported^22^, the other NDP components show resistance to gene amplification, with no more than four copies per genome (**Fig. 1a**). As central hubs for cellular housekeeping and plant development, MetAP, ATE, PRT6 and BIG have ben shown to function within complexes in animal cells^38^, thus requiring a finely tuned stoichiometry between subunits also in plants, as highlighted by our ERC analysis (**Fig. 4**). Consequently, plants are likely sensitive to altered dosage of these proteins, constraining their duplication and diversification.

PCOs and ERFVIIs have undergone greater expansion and diversification, although with constraints reflected in features conserved across all angiosperms analyzed in this study, regardless of the probability of submergence in their original habitats (**Fig. 1**). Notably, among the most conserved feature of ERFVII, we found the Cys-degron, indicating the requirement to subjugate stability of these proteins to conditions that reduce O_2_ or NO levels^39,40^. Such conditions are likely driven by endogenous processes, including developmental and growth programs^9,41^. This interpretation is supported by our oxygen-profiling survey, which revealed distinct oxic states between differentiated and meristematic tissues in nearly all species examined (**Fig. 2a**). An unexpected exception was observed in *C. gigantea*, which does not maintain their SAM at reduced oxygen tension but nevertheless retain CMVII-1 in their ERFVIIs (**Fig. 2a**). This finding suggests the presence of other hypoxic tissues or cell types that require ERFVII-mediated transcriptional regulation. Interestingly, *C. gigantea* was the only species among the six we investigated whose hypoxia-responsive genes did not show a significant enrichment of HRPEs in their promoters (**Fig. 5f**). It could be speculated that the absence of developmental hypoxia in the SAM may have reduced the selective pressure to maintain ERFVII dominance over hypoxia-responsive genes.

Adaptation to aquatic environments in Ceratophyllales and monocots correlates with an increase in ERFVII genes and a reduction of A-type PCO sequences. This trend, already highlighted from the study of seagrass genomes^18^, is particularly evident in monocots, where the two Cys-NDP components show the most striking variations in aquatic species. Within the PCO orthogroup, A-type genes were entirely lost, while C-type genes expanded, with some acquiring hypoxia inducibility (**Fig. 1a, Fig. 5b**) and others potentially evolving a Cys-degron (**Supplementary Fig. 2d**). Similarly, ERFVIIs underwent dramatic expansion in aquatic monocots, including the emergence of a novel clade characterized by a distinct domain architecture, termed HRE_aqua_ (**Fig. 1c**). At the same time, this clade shows recurrent loss of the Cys-degron or mutations in its consensus sequence, abolishing participation as a Cys-NDP substrate (**Fig. 3**). The impact of aquatic habitats on PCOs and HREs, together with their feedback regulation in response to O_2_ availability, suggests a potential co-dependent evolution of these two Cys-NDP components. A similar interdependence was recently observed when reconstituting the Cys-NDP module for hypoxic gene induction in yeast, where interlocked feedback loops were required to ensure sustained and rapid responses^42^.

The evolutionary connection between PCO and ERFVII extends beyond feedback regulation. While N-terminal cysteine oxidases likely originated in the last common ancestor of eukaryotes^43^, B-type PCOs emerged in vascular plants, concomitant with the recruitment of ERFVII as transcriptional mediators of low-oxygen signaling^1,22^. We speculate that constitutively expressed A-type PCOs may have become dispensable in aquatic species that regularly encounter nocturnal hypoxia due to submergence. Alternatively, it may not be constant hypoxia but rather extreme O_2_ fluctuations, caused by intertidal habitats or reduced gas diffusion in water, that shaped this trajectory, with environments becoming hyperoxygenated in the light and severely O_2_-depleted at night.

Our survey did not only look at the components of the Cys-NDP but also considered the transcriptional output of low oxygen signaling. We identified a conserved core of 11 upregulated orthogroups (OGs), including genes encoding fermentation and signaling proteins, most of which are regulated by ERFVII in *Arabidopsis*^1^ (**Fig. 5**). Many of these OGs are also induced by hypoxia in non-flowering vascular plants^44^. For some OGs, such as those containing fermentation genes, hypoxia induction is also observed in non-vascular species that lack ERFVIIs, indicating that this response was already present in the last common ancestor of land plants^1,45^. Collectively, these observations suggest that ERFVIIs were recruited early to regulate these core response genes, enabling tighter, or possibly stronger, induction under hypoxia. The conservation of this response across all vascular plants highlights its necessity for maintaining NAD+ regeneration or pyruvate consumption when ATP synthesis is restricted to glycolysis^46^.

Our analysis also confirmed the conserved induction of the trihelix HRA1, LBD factors, and MYBs in angiosperms. The first was shown to be required to attenuate gene induction in young tissues in Arabidopsis and the presence of a repressive motif in the second suggests a similar function^12,47,48^. The MYB11/11/112 clade, instead, has been characterized as a positive regulator of flavonoid synthesis in Arabidopsis^49^, compounds that have been shown to accumulate in different species in response to submergence and waterlogging^50–52^. Their inactivation led to somehow enhanced susceptibility to strict hypoxia (**Fig. 5d**), although the effect is not as dramatic as in the case of fermentation mutants^45,53^. The consistent induction of these genes across angiosperms, regardless of their likelihood of experiencing ambient hypoxia, suggests they are primarily required for developmental or environmental responses involving hypoxia, rather than specifically for flooding tolerance. For instance, LBD40/41 was proposed to regulate somatic embryogenesis^36^, while MYB11/111/112 modulate flavonoid biosynthesis in response to diverse abiotic stresses^49^, likely contributing to reactive oxygen species (ROS) homeostasis^54^.

When examining potential habitat-driven influences on hypoxia responses, we found a surprising pattern: although aquatic plants show similar trends in the composition of the Cys-NDP (**Fig. 1**), the two species analysed in our transcriptomic survey, *C. demersum* and *Z. marina*, shared few DEOGs (**Fig. 5b**). This is partly explained by their attenuated hypoxia responses, but it is also possible that ERFVIIs recruited different gene sets to accommodate hypoxia responses tailored to their distinct anatomies. By contrast, epiphytes and xerophytes shared several DEOGs (**Fig. 5b**). We speculate that the same endogenous selective pressures that favour retention of Cys-NDP components in the absence of submergence may also shape the repertoire of hypoxia-response genes. Such regulation could be required to support metabolism or differentiation in specific tissues or cell types.

In conclusion, we carried out a pioneering analysis of the genetic diversity available among plants whose genome has been sequenced, searching for features that could connect submergence with the know mechanism for oxygen sensing. We show that, counterintuitively, reduced chances of environmental hypoxia do not alleviate selective pressure to retain the core the oxygen sensing machinery. We hypothesized that this is linked to anatomical or metabolic requirements to maintain subjugation to the NDP to cope with oxygen fluctuations, such as cyclic hypoxia^41^. Instead, persistent submergence favored expansion and diversification of the hypoxia regulators and allowed their release from Cys-NDP-dependent regulation. As sequencing technologies become cheaper, data for more species from these habitats will become available allowing re-assessment of these hypotheses. They will also allow the identification of more exceptions to the conserved oxygen sensing mechanism which provide precious information for breeders and molecular physiologists for the engineering of genetic circuits aimed at crop improvement.

## Methods

### Plant material and growth conditions

Seven species were analyzed: *Ceratophyllum demersum* (Ceratophyllaceae), *Zostera marina* (Zosteraceae), *Tillandsia fasciculata* (Bromeliaceae), *Dendrobium nobile* (Orchidaceae), *Kalanchoë fedtschenkoi* (Crassulaceae), *Viscum album* (Santalaceae) and *Carnegiea gigantea* (Cactaceae).

Aquatic species (*C. demersum* and *Z. marina*) were maintained in 50 L tanks at 22 °C under long-day conditions (16 h light/8 h dark, 50–60 μmol m−^2^ s−^1^ white light). *C. demersum* plants were obtained from Pond Plant Growers Direct (Burgh-Le-Marsh, UK). *Z. marina* seeds were collected from Orkney, Scotland (58°55′13.0″ N, 2°47′51.0″ W; 58.920278, –2.797500), stratified for 30 d at 4 °C in artificial seawater at 25‰, germinated in artificial seawater at 10‰, and subsequently transferred to 25‰ for growth until the emergence of three true leaves. Tanks were continuously aerated.

Terrestrial species were cultivated in controlled-environment growth chambers at either 22 °C (*T. fasciculata, D. nobile, V. album*) or 24 °C (*K. fedtschenkoi, C. gigantea*). *T. fasciculata* plants were obtained from GrowTropicals (Leeds, UK), *D. nobile* from Aylett Nurseries (St Albans, UK), *K. fedtschenkoi* from Allwoods (Hassocks, UK), and *C. gigantea* seeds from Pretty Wild Seeds (Darlington, UK). *V. album* shoots were collected from the Private Fellows’ Garden in Wadham College, Oxford, UK. All species were maintained under long-day conditions (16 h light/8 h dark, 50–60 μmol m−^2^ s−^1^ white light). *K. fedtschenkoi* and *C. gigantea* were grown in a 1:1:1 mixture of peat-free compost, sand and vermiculite; *C. gigantea* seedlings were germinated in peat-free compost. *D. nobile* was cultivated in a peat-free orchid substrate consisting of a 7:2:1 mixture of pine bark, perlite and Seramis clay.

*Arabidopsis thaliana* ecotype Columbia-0 (Col-0) was used as the wild type. The *myb11/12/111* triple mutant^49^ (N9815, CS9815) was obtained from the Nottingham Arabidopsis Stock Centre (NASC). Seeds were surface-sterilized and stratified at 4 °C for 3 d in the dark before germination. For in vitro propagation, seeds were cultivated on half-strength Murashige and Skoog (MS) medium (Duchefa) supplemented with 1% (w/v) sucrose and 0.8% (w/v) agar. Plants were grown at 22 °C/20 °C (day/night) under a 16 h light/8 h dark photoperiod with a light intensity of 100 μmol m−^2^ s−^1^. Plates were positioned vertically after stratification to allow seedling development.

### Generation of the Arabidopsis *lbd40/41/42* mutant using CRISPR

Guide RNAs (gRNAs) targeting *LBD40* (*At1G67100*) and *LBD41* (*At3G02550*) were cloned into the binary plasmid pKSE401, and a gRNA targeting *LBD42* (*At1G68510*) was cloned into pHSE401 as described by Xing *et al*. (2014) ^55^. The resulting plasmids were used for *Agrobacterium*-mediated transformation of Columbia-0 plants through successive rounds of floral dip infiltration. First-generation transformant (T1) plants were selected on antibiotics (hygromycin for pHSE401 and kanamycin for pKSE401). The Cas9-bearing cassette was subsequently removed by segregation in later generations. A plant carrying frame-disrupting mutations in each *LBD* gene was identified using PCR amplification with gene-specific primers followed by Sanger sequencing (**Supplementary Fig. 8**). Seeds from this triple mutant were collected for subsequent experiments.

### Comparative genomics

Orthogroups and the species tree were inferred with OrthoFinder v2.5.5^21^ using protein-coding primary transcript sequences from 54 angiosperm species, representing *Nymphaeales* (1), *Ceratophyllales* (1), monocots (24) and eudicots (28) (Supplementary Table X). *Marchantia polymorpha* and *Physcomitrium patens* were included as outgroups (**Supplementary Table 1**). OrthoFinder was run with default parameters using 96 parallel threads. The OrthoFinder species tree was examined, confirmed to be consistent with established angiosperm phylogenies, and used for all downstream analyses.

ERC analyses were performed using ERCnet^30^ applied to the OrthoFinder output. An all-versus-all ERC calculation was conducted with parameters -s -e -p 12 -r 10 -n 2 -T. To specifically assess the N-cys degron components, ERCnet was run with parameters -e -s -p 12 -r 10 -l 100 -n 2 -a -T. Significant ERC pairs were defined by *P* ≤ 0.05 and R^2^ ≥ 0.2 in both Spearman and Pearson correlation, and downstream functional analyses were restricted to ERC pairs meeting these criteria.

### Motif signature analysis

Motif discovery was carried out with MEME Suite v5.4^56^ using default parameters to annotate conserved amino acid motifs in ERFVII proteins.

### Phylogenetic analysis

Protein sequences of ERFVII orthologs from the sampled angiosperms were aligned with MAFFT v7.505^57^ using the L-INS-i algorithm (--localpair --maxiterate 1000) and 48 threads. Maximum-likelihood phylogenetic analyses were performed with IQ-TREE v2.2.2.7^58^under the best-fit substitution model selected by ModelFinder (JTT+R8). Branch supports were estimated using 1,000 ultrafast bootstrap replicates and SH-aLRT tests. Phylogenetic trees were visualized and annotated with Interactive Tree of Life (iTOL)^59^.

### 3D structure prediction

The crystal structure of AtPCO4 was retrieved from the Protein Data Bank (PDB ID: 6S7E). Predicted structures of consensus type-C PCOs were generated using AlphaFold2^60^. All structures were visualized and aligned using PyMOL^61^.

### Hypoxia treatments

For aquatic species, whole plants of *C. demersum* and plantlets of *Z. marina* with three true leaves were exposed to hypoxia by bubbling 1% (v/v) O_2_ in N_2_ for 8 h in the dark (00:00–08:00). Control plants were maintained under the same conditions while bubbling atmospheric air (21% O_2_). For terrestrial species, whole adult plants of *T. fasciculata, D. nobile* and *K. fedtschenkoi*, shoots of *V. album*, and 1-month-old plantlets of *Carnegiea gigantea* were exposed to 1% (v/v) O_2_ in N_2_ for 8 h in the dark (00:00–08:00) using the H35 Hypoxistation (Don Whitley Scientific). Control plants were maintained in the dark under atmospheric air (21% O_2_). Each biological replicate consisted of three individuals per species.

*Arabidopsis thaliana* Col-0, *myb11/12/111* and *lbd40/41/42* mutants were grown on vertical plates and subsequently exposed to either air (21% O_2_) or severe hypoxia (0.1% O_2_) for 24 h in a H35 Hypoxystation (Don Whitley Scientific). During treatments, plates were kept in darkness to prevent O_2_ release by photosynthesis. Following hypoxia, plates were returned to air and maintained under normoxic conditions for 5 d, after which fresh biomass and survival were assessed.

### RNA isolation and sequencing

Total RNA was isolated from 80–100 mg of frozen, ground young shoots of *Tillandsia fasciculata, Dendrobium nobile, Kalanchoë fedtschenkoi, Viscum album* and *Ceratophyllum demersum*, and from photosynthetic tissues of *Zostera marina* and *Carnegiea gigantea*, using the GeneJET Plant RNA Purification Kit (Thermo Fisher Scientific) according to the manufacturer’s instructions. RNA quantity and purity were assessed by spectrophotometric measurements of OD_260_/OD_280_ (NanoDrop 330, Thermo Fisher Scientific) and by 2% (w/v) agarose gel electrophoresis. RNA integrity was verified with RNA Integrity Number (RIN) values, and samples with RIN > 7.5 were used for cDNA library construction. Libraries were prepared and sequenced at Novogene Bioinformatics Technology Co., Ltd. (Beijing, China) on an Illumina NovaSeq 6000 platform, generating 2 × 150 bp paired-end reads.

### Differential gene expression analysis

RNA-seq libraries were mapped to their respective reference genomes (Supplementary Table X) as described by Althiab-Almasaud et al. (2021)^62^. Differential expression analysis was performed in R v4.4.1 (https://www.r-project.org/) using DESeq2 v1.46^63^ with default relative log expression (RLE) normalization. The false discovery rate (FDR) was controlled using the Benjamini–Hochberg method, and genes were considered differentially expressed (DEGs) at adjusted *P* (padj) < 0.05 and log_2_FC > |1|. Because RLE normalization in DESeq2 does not account for transcript length, normalized counts were further corrected by transcript size and read length to allow expression comparisons in downstream analyses such as profile clustering. Specifically, DESeq2-normalized counts were divided by transcript length and multiplied by the raw read length, providing relative estimates of transcript numbers.

### Microprofiling of O_2_ in the SAM

Microprofiling of tissue O_2_ followed a previously published procedure^9^ with modifications. Young shoots of *C. demersum, Z. marina, T. fasciculata, D. nobile, K. fedtschenkoi, V. album* and *C. gigantea* were sectioned to expose the shoot apical meristem (SAM) and secured in Petri dishes containing 8% (v/v) agar. O_2_ microprofiles in the SAM were obtained using a custom-built Clark-type microsensor with a bevelled 3 μm tip (Unisense A/S), connected to a picoammeter (Oxymeter, Unisense A/S) and mounted on a motorized micromanipulator (MM33, Unisense A/S). Data acquisition and manipulator positioning were controlled with SensorTrace Suite v2.8 (Unisense A/S). The microsensor tip was advanced in 20 μm steps, starting outside the tissue and continuing until the target tissue was fully penetrated.

### Vibratome sectioning and confocal microscopy

After O_2_ measurements, samples were fixed in 70% ethanol:acetic acid (3:1) for 24 h and then transferred to 70% ethanol. Fixed samples were washed twice in 1× PBS, embedded in 6% agarose (in 1× PBS), and sectioned into 50 µm cross-sections using a vibratome. Sections were stained in 1× PBS supplemented with Renaissance SCRI 2200 (SR2200; 1 µl ml−^1^; Renaissance Chemicals) to label cell walls^64^. Prior to imaging, sections were washed twice with 1× PBS.

Confocal imaging was performed with a Leica Stellaris confocal laser scanning microscope (Leica Microsystems). Images were stitched using LAS X Life Science Microscope Software, and the shoot apical meristem (SAM) was identified across serial sections.

### Yeast DLOR assay

AtPCO4 in pAG415GPD and C-DLOR in pAG413GPD were obtained from Lavilla-Puerta et al. (2023)^29^. DLOR versions comprising the first 50 aa of the selected ERFVIIs (Supplementary Table 5) were synthesized and cloned in pENTR221 by GeneArt (Thermo Fisher Scientific) and recombined into pAG413GPD (Addgene plasmid #14142) via Gateway LR® clonase, following the manufacturer’s protocols (Thermo Fisher Scientific). Each DLOR version was co-transformed with AtPCO4-pAG415GPD into BY4742 yeast (*Matα; his3-Δ1; leu2-Δ0; lys2-Δ0; ura3-Δ0*). Yeast cells were selected and grown in synthetic drop-out (SD) media without histidine and leucine (SD -leu -his) as described previously (10.1016/j.jmb.2019.05.023). Up to 6 independent yeast colonies per genotype were diluted to an OD600 of 0.1 in a flat-shaped 96-well microplates, in 200 µl of SD -leu -his and kept for 6 h at either 1% (hypoxia) or 21% (aerobic control) v/v O2 at 30 °C. Plates were then spun for 5 min at 2000 rpm and pellets frozen at -70 °C. Frozen yeast pellets were resuspended in 50 µl of 1x PLB, and 6 µl of the lysate were used to measure Firefly and Renilla luciferase (normalized in Fluc/Rluc) using the Dual-Luciferase Reporter (DLR) Assay System (Promega) following the manufacturer’s instructions.

## Data availability

RNA sequencing raw data generated for this study was deposited in the Sequence Read Archive (SRA) at the National Centre for Biotechnology Information under BioProject ID SUB15641052.

## Use of Large Language Models (LLMs)

ChatGPT 5 was used to help polish or refine the text of this manuscript.

## Acknowledgements

The authors would like to thank Evan S. Forsythe (Oregon State University) for extensive discussion and advice on the application of the ERC analysis Chiara Perico (University of Oxford) for assistance with tissue sectioning and advice on SAM imaging. Judith Bäumler and Marvin Hönle (University of Bayreuth) are acknowledged for initial work on the *lbd* mutant lines. This work was supported by the European Research Council (ERC) grant 101001320 (FL and VS) and by the UKRI Biotechnology and Biological Science Research Council (BBSRC) grants BB/X001059/1 (FL and XC), CBR00770 (FL and ML-P) and CBR01420 (FL and VS).

## Supplementary Figures

**Supplementary Figure 1**. ERFVII evolution across the ecological/functional categories of plants, based on their habitat adaptations.

**a**. Bubble plot depicting the distribution of RAP, HRE and non-MC ERFVIIs across macrophytic, epiphytic, xerophytic and mesophytic angiosperms. **b**. Box plot depicting the ratios of ERFVII-to-PCO gene numbers in macrophyte, xerophyte, epiphyte and mesophyte angiosperms. **c**. Number of non-MC ERFVIIs across macrophytic, epiphytic, xerophytic and mesophytic angiosperms.

**Supplementary Fig. 2**. The C-group PCOs.

**a**. Multialignment of logos representing the variation of residues in conserved regions of type A, B and C PCOs. **b**. Alpha-fold predicted structure (ribbon) of the consensus of type-C PCOs. The His residues predicted to be involved in FeII coordination as well as other conserved residues that are relevant for enzyme catalysis are shown as balls and sticks. **c**. Surface of the predicted sequence shown in c, showing the predicted substrate cavity. **d**. Multialignment of the N-termini of three type C PCOs from sea grasses (*Zostera marina, Zostera muelleri* and *Potamogeton acutifolius*).

**Supplementary Fig. 4**. Oxygen profiles measured across tissues and stems of the species analysed in Fig. 2.

**Supplementary Fig. 5**. Oxygen levels measured in aquaria.

a. A scheme of the experimental setup b. Oxygen concentrations as measured by a clarck electrode placed in the aquaria close to the plants.

**Supplementary Fig. 6**. Differentially expressed genes of *Zostera marina, Ceratophyllum demersum, Kalanchoë fedtschenkoi, Carnegiea gigantea, Viscum album, Dendrobium nobile* and *Tillandsia fasciculata*.

Volcano plots showing relative expression levels between hypoxic and aerobic plants (log_2_ fold change) and statistical significance (-log_10_p. adj.). Purple and green dots indicate significantly up- and down-regulated genes, respectively.

**Supplementary Fig. 7**. Gene expression changed caused by hypoxia in A. thaliana for LDB and MYB genes analysed in Fig. 5c-e.

**Supplementary Fig. 8. Generation of triple *lbd* knock-out mutant. a**. List of priemrs used to create the gRNAs for gene knock-out. **b**. Scheme depicting frame shift created by CRISPR in Arabidopsis *LBD40, LBD41* and *LBD42*.

## Supplementary Tables

**Supplementary Table 1 | Distribution of Cys-NDP components across 54 angiosperms**.

Species metadata (clade, order, habitat, taxonomy, genome reference) and the number of genes per orthogroup are shown, including ERFVII subtypes (HRE, No-MC_ERFVII, RAP), MC-ZPR2, VRN2, PCO, ATE, PRT6, BIG, AtMetAP2The ERFVII/PCO gene ratio is provided for each species.

**Supplementary Table 2 | Motif analysis of ERFVII transcription factors in 54 angiosperms**.

Sheet 1 lists the motifs identified by MEME, including the motif ID, consensus sequence, motif width (amino acids), and E-value.

**Supplementary Table 3 | Evolutionary rate covariation (ERC) analysis of Cys-NDP components**.

Each row reports a pairwise ERC comparison between GeneA and GeneB, with their corresponding orthogroup identifiers (HOG). The number of overlapping branches used for the ERC calculation is shown, together with Pearson’s correlation coefficient (P_R^2^) and significance (P_Pval), and Spearman’s correlation coefficient (S_R^2^) and significance (S_Pval). Columns Interactome, MC, and MC9 indicate whether the gene pair is supported by known interaction data or associated with the metacaspase (MC9) or proteins starting by MC.

**Supplementary Table 4 | Transcriptomic response of selected epiphytes (D. nobile, T. fasciculata and V. album), macrophytes (*C. demersum* and *Z. marina*), xerophytes (C. *gigantea* and *K. fedtshenkoi*) and mesophytes (*A. thaliana*) to hypoxia**.

For all species, except Arabidopsis, two gene IDs are shown, followed by log2fold change, adjusted p-value, HOG and OG.x

**Supplementary Table 5 | Cys-N-degron variants of selected species**.

The protein sequences are shown first, followed by the DNA sequences considered for cloning.

