## Supplementary Figures for "Anatomy and habitat shape the oxygen sensing machinery of angiosperms"

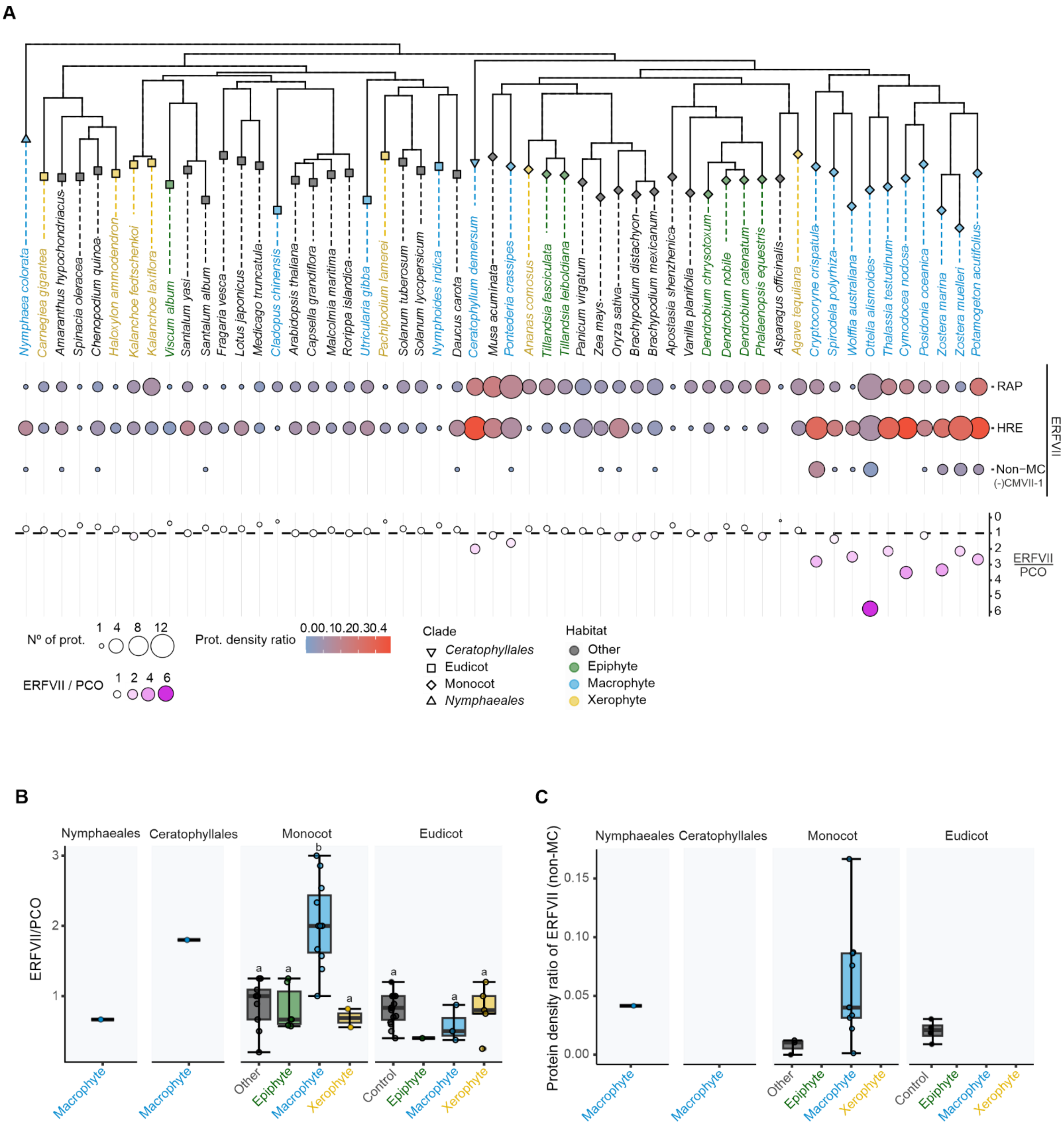


**Supplementary Figure 1.** ERFVII evolution across the ecological/functional categories of plants, based on their habitat adaptations.

**a.** Bubble plot depicting the distribution of RAP, HRE and non-MC ERFVIIs across macrophytic, epiphytic, xerophytic and mesophytic angiosperms. **b.** Box plot depicting the ratios of ERFVII-to-PCO gene numbers in macrophyte, xerophyte, epiphyte and mesophyte angiosperms. **c.** Number of non-MC ERFVIIs across macrophytic, epiphytic, xerophytic and mesophytic angiosperms.

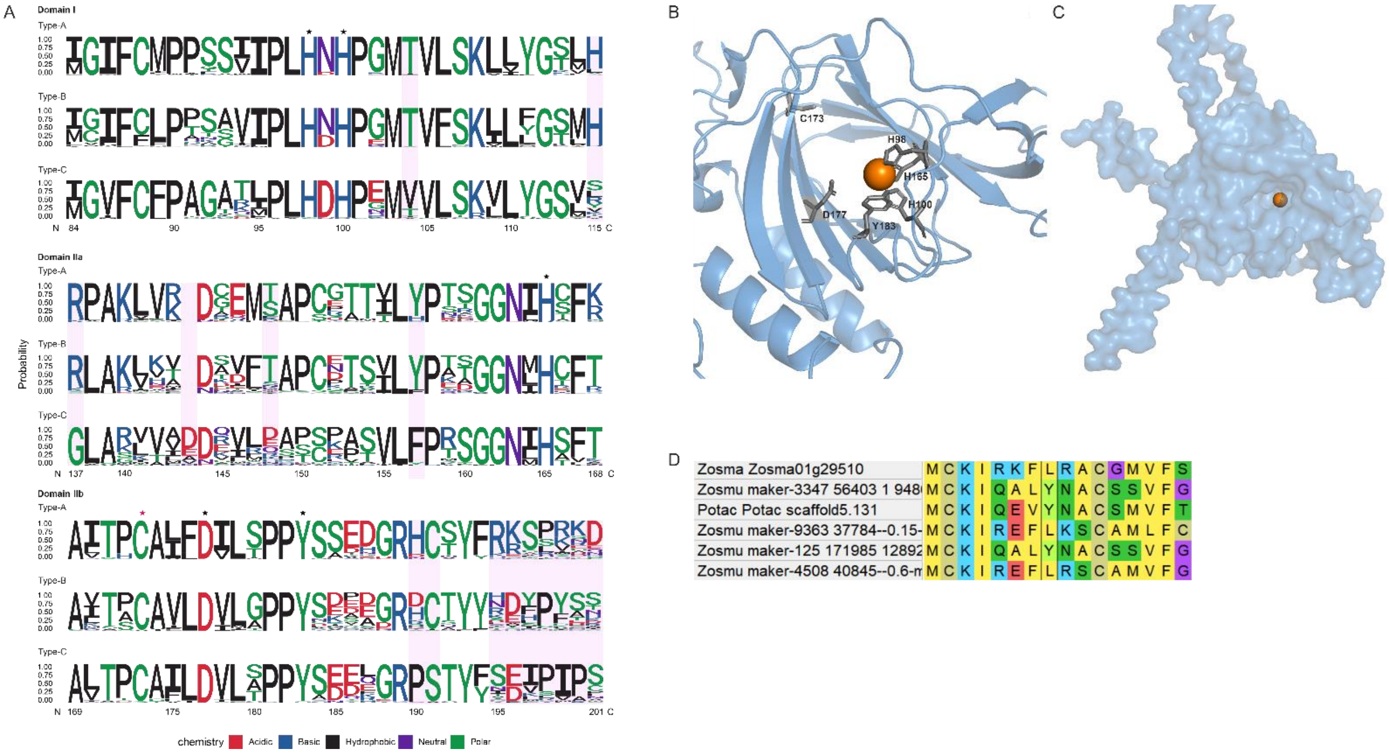


**Supplementary Fig. 2.** The C-group PCOs.

**a.** Multialignment of logos representing the variation of residues in conserved regions of type A, B and C PCOs. **b.** Alpha-fold predicted structure (ribbon) of the consensus of type-C PCOs. The His residues predicted to be involved in FeII coordination as well as other conserved residues that are relevant for enzyme catalysis are shown as balls and sticks. **c.** Surface of the predicted sequence shown in c, showing the predicted substrate cavity. **d.** Multialignment of the N-termini of three type C PCOs from sea grasses (*Zostera marina*, *Zostera muelleri* and *Potamogeton acutifolius*).

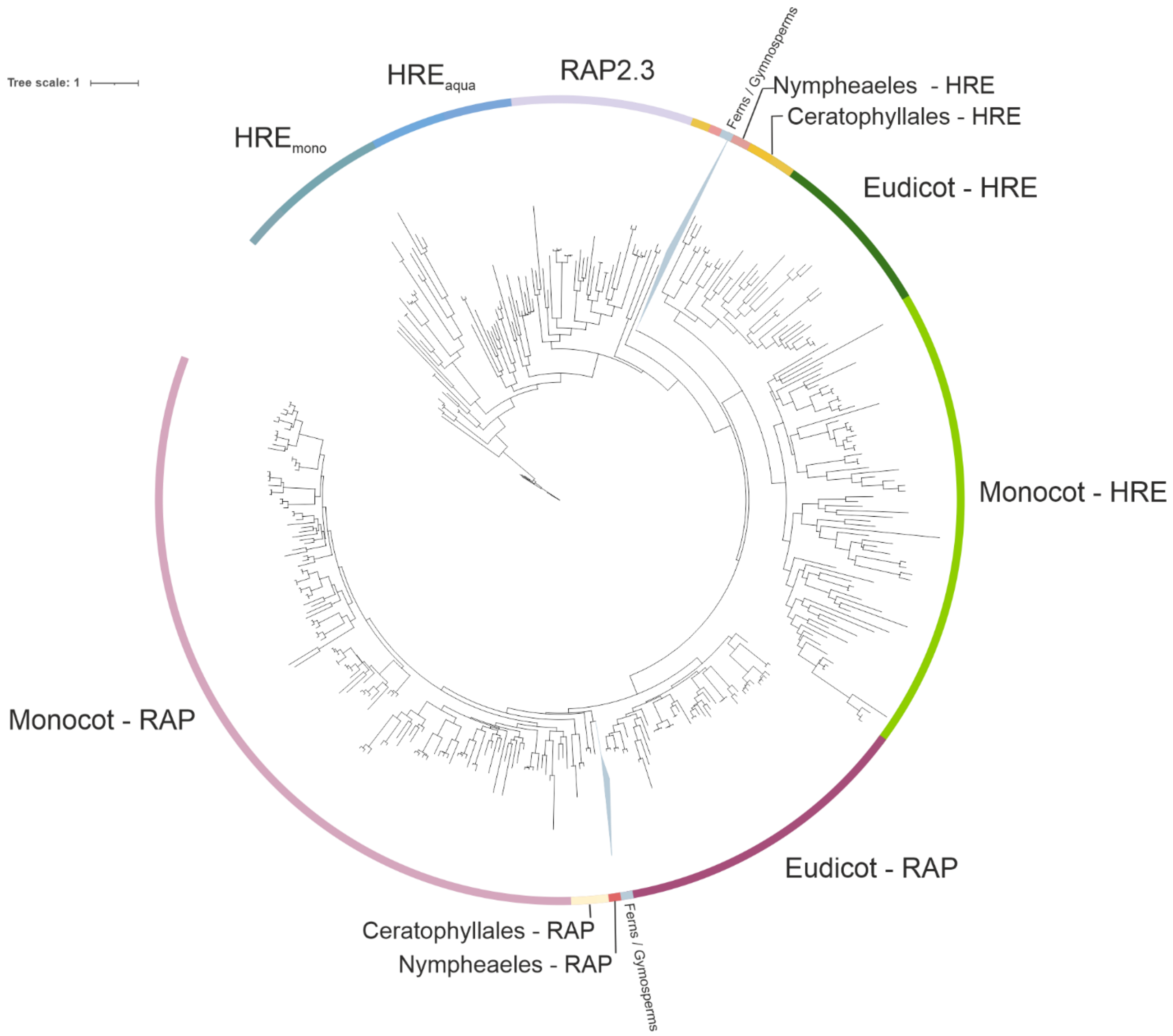


**Supplementary Figure 3**. Phylogenetic tree illustrating the relatedness of group VII ERFs among the 54 plant species considered in Figure 1.

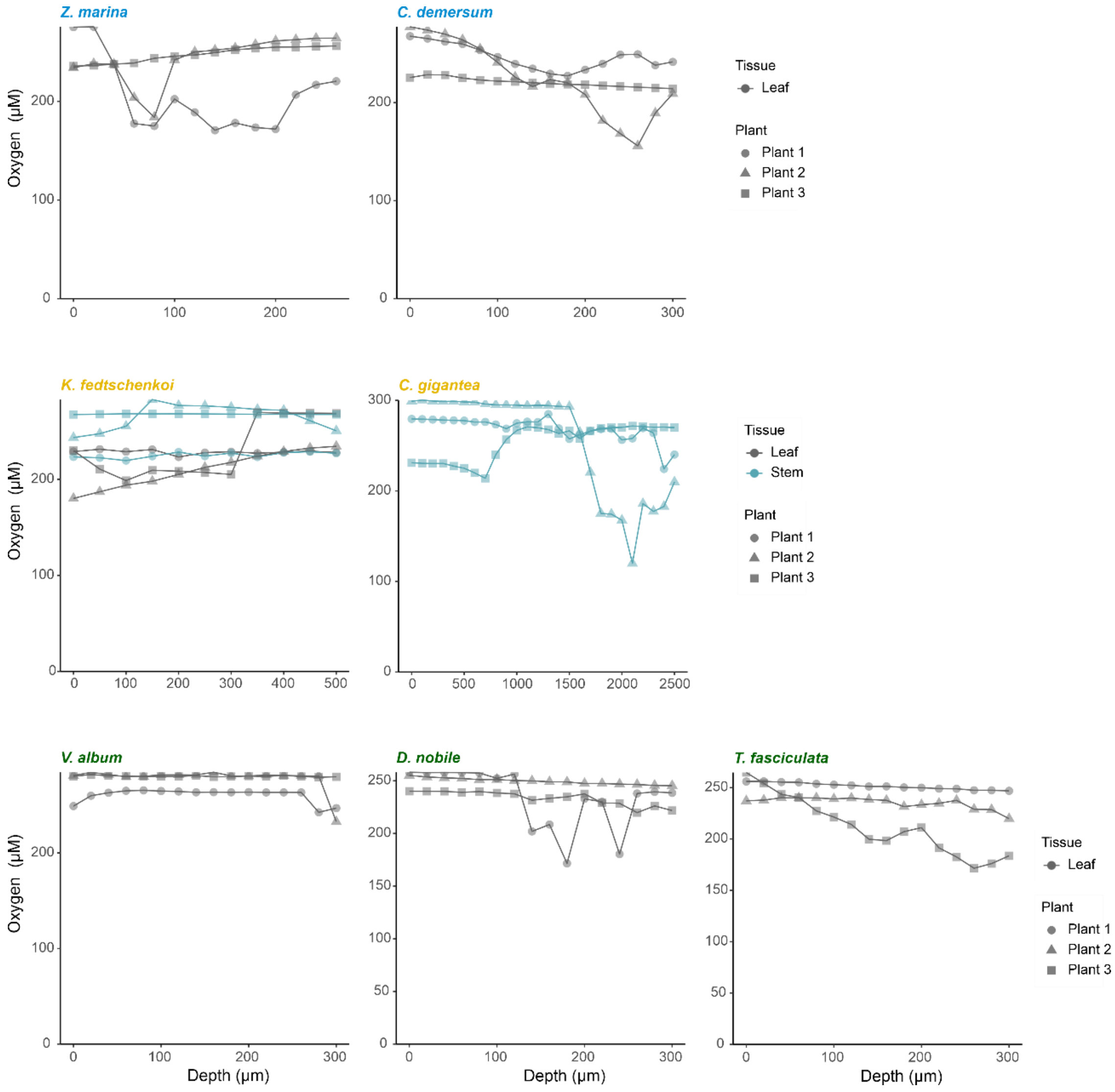


**Supplementary Fig. 4.** Oxygen profiles measured across tissues and stems of the species analysed in Fig. 2.

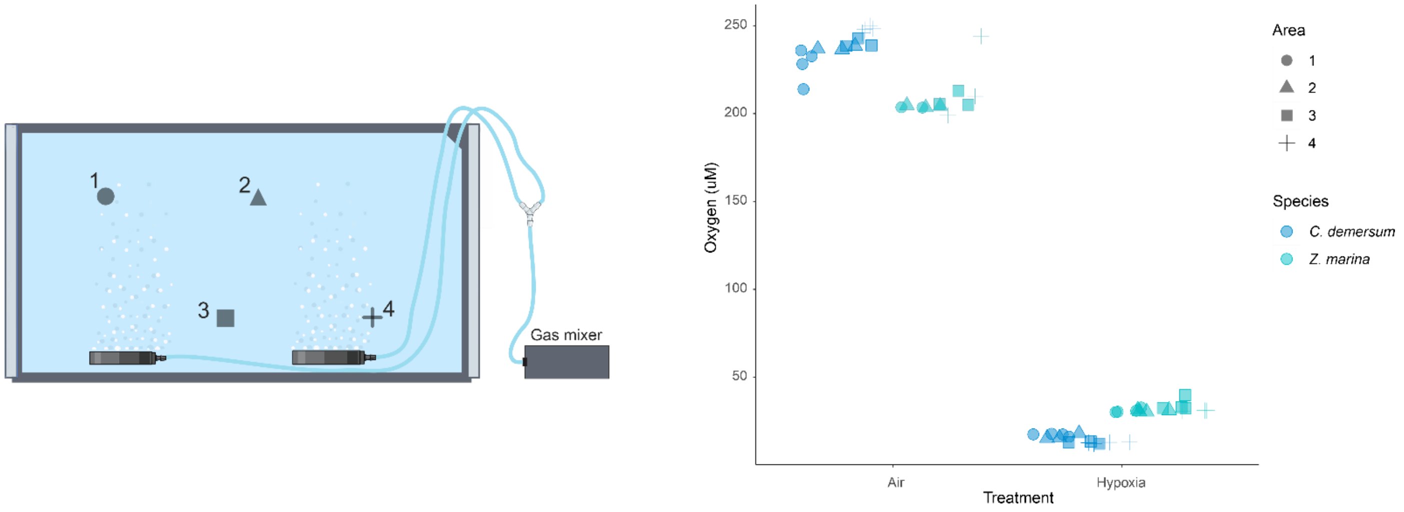


**Supplementary Fig. 5.** Oxygen levels measured in aquaria.
a. A scheme of the experimental setup b. Oxygen concentrations as measured by a clarck electrode placed in the aquaria close to the plants.

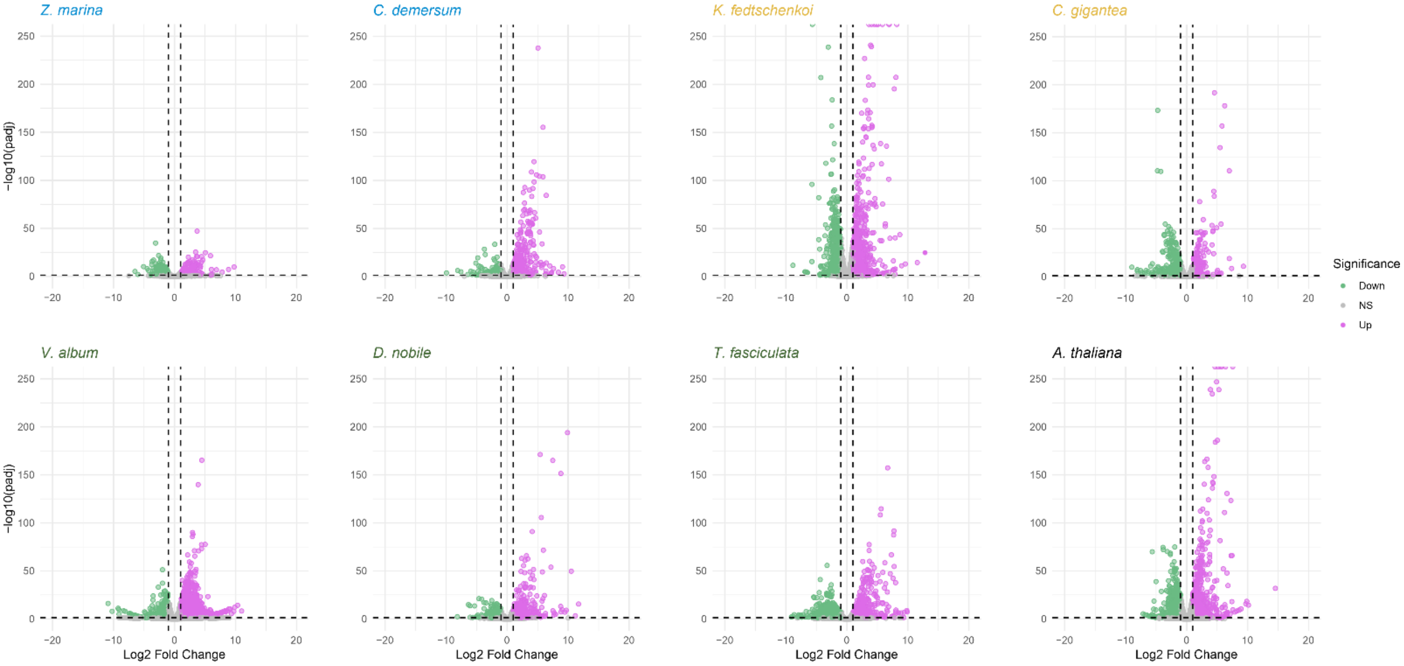


**Supplementary Fig. 6.** Differentially expressed genes of *Zostera marina*, *Ceratophyllum demersum*, *Kalanchoë fedtschenkoi*, *Carnegiea gigantea*, *Viscum album*, *Dendrobium nobile* and *Tillandsia fasciculata*.
Volcano plots showing relative expression levels between hypoxic and aerobic plants ($\log_2$ fold change) and statistical significance (-$\log_{10}$p. adj.). Purple and green dots indicate significantly up- and down-regulated genes, respectively.

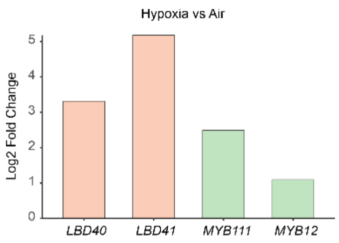


**Supplementary Fig. 7.** Gene expression changed caused by hypoxia in A. thaliana for LDB and MYB genes analysed in Fig. 5c-e.

**A**

| Primer name | Primer sequence | Vector |
|---|---|---|
| AtLBD41_Crispr1_fw | ATTGAAAACCACTAACCAGGACGA | pKSE401 |
| AtLBD41_Crispr1_rev | AAACTCGTCCTGGTTAGTGGTTTT | |
| CRISPR_LBD40D_fw | ATTGACCGCTGCGTCGCATGCGAT | pKSE401 |
| CRISPR_LBD40D_rev | AAACATCGCATGCGACGCAGCGGT | |
| LBD42_CRISPR2_fw | ATTGGGTGCGACGCGGATGCACT | pHSE401 |
| LBD42_CRISPR2_rev | AAACAGTGCATCCGCGTCGCACC | |

**B**

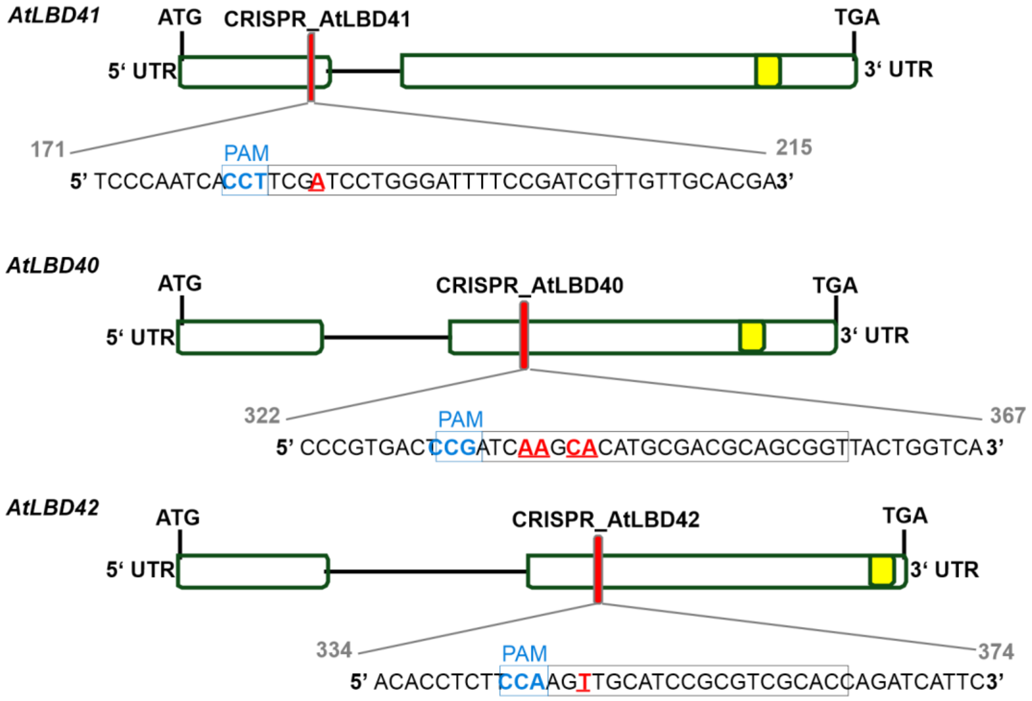


**Supplementary Fig. 8. Generation of triple *lbd* knock-out mutant.**
**a.** List of priemrs used to create the gRNAs for gene knock-out. **b.** Scheme depicting frame shift created by CRISPR in Arabidopsis *LBD40*, *LBD41* and *LBD42*.
